# AlphaRING-X: accurate interpretation of missense variant deleteriousness based on protein structural stability

**DOI:** 10.1101/2024.11.12.623182

**Authors:** Aaron Mamane-Logsdon, Roghaiyeh Safari, Anna Barnard, Goedele N. Maertens, Evangelos Bellos

## Abstract

**Background:** Accurate interpretation of missense variants remains a significant challenge hindering genomic diagnosis. While state-of-the-art machine learning and deep learning predictors offer high accuracy, they often lack the transparency required for clinical evidence weighting. To bridge this gap between accuracy and clinical utility, we developed AlphaRING-X.

**Methods:** AlphaRING-X is an open-source framework that integrates local and global protein structural stability features using an efficient implementation of gradient boosting, XGBoost, to predict variant deleteriousness. Local stability is interrogated through a combination of AlphaFold-based modelling and residue interaction network analysis. Changes in global stability upon mutation (ΔΔG) are estimated using FoldX. For each prediction, AlphaRING-X quantifies the magnitude and direction of each feature’s contribution using Shapley additive explanations (SHAP), providing a unified yet interpretable score. We trained and evaluated the model using a gold standard ClinVar dataset of validated neutral and deleterious variants.

**Results:** AlphaRING-X achieved an area under the receiver operator curve of 0.94, significantly outperforming widely used predictors such as CADD. Crucially, it provided unambiguous classification for 95.5% of variants at 90% precision in both neutral and deleterious classes. We analysed each prediction’s SHAP values, identifying local connectivity and disorder as the strongest contributors.

**Conclusions:** AlphaRING-X combines high-performance prediction with the interpretability necessary for clinical decision-making. Its flexible, open-source architecture makes it a powerful, transparent tool for enhancing genomic diagnostics and advancing precision medicine.

## Background

Missense variants dominate the landscape of human protein-coding variation, with a recent catalogue of one million exomes identifying over 10.4 million unique missense variants at a median burden of 8,700 per individual [1]. Such variants alter protein sequence and, therefore, function to varying degrees, producing phenotypic diversity. This ranges from benign variation to Mendelian disorders and measurable risk for complex diseases [1,2,3]. Therefore, accurate interpretation of the deleteriousness of missense variants is essential to determine which can be considered diagnostic and thus actionable for the clinician and returnable to the patient.

Guidelines have been established to assess variant actionability, most notably by the American College of Medical Genetics and Genomics and the Association for Molecular Pathology (ACMG/AMP), which states that after considering population, computational, predictive, functional, segregation, de novo, allelic, and other annotations, there must be at least 90% certainty in a variant’s deleteriousness for a classification to be made [4]. To facilitate access to variant classifications, multiple databases have been curated to catalogue available variant annotations, including general-purpose resources such as ClinGen [5,6] and ClinVar [7,8], as well as specialised resources such as COSMIC [9]. However, definitive classification remains unavailable for the majority of variants. This gap largely stems from the difficulty of gathering the diverse, orthogonal lines of annotation required by evidence-weighting frameworks like ACMG/AMP for variants that are inherently rare. As a result, it has been estimated that only 2% of known missense variants have a classification [10].

To alleviate this burden, many computational tools have been developed to readily predict variant deleteriousness, thereby allowing researchers and clinicians to investigate further those with a greater likelihood of being actionable. Early efforts, such as SIFT [11] and PolyPhen [12], were based on evolutionary constraints, which could effectively capture the deleteriousness signal. However, these tools relied heavily on phylogenetic conservation, meaning they struggled with poorly aligned, fast-evolving, and lineage-specific regions of proteins. A significant advancement was the development of CADD [13], which has undergone continuous improvement over the last decade [14,15,16]. CADD uses a logistic regression model trained using hundreds of DNA-, RNA-, and protein-level features to output genome-wide PHRED-scaled deleteriousness scores for variants [16]. Although CADD’s extensive and varied feature space ensures accurate predictions across diverse variants, the contributions of these features to deleteriousness scores are not reported, hindering mechanistic interpretation and downstream clinical utility. Similarly, recently developed deep learning-based tools [17,18,19,20,21] are accurate but, due to their black-box architectures, struggle to explain the gap between their outputs and the underlying biophysical mechanisms driving the variant’s effect.

To address the above challenges, we aimed to develop a framework for missense variant deleteriousness prediction that is both highly accurate and easily interpretable, resulting in AlphaRING-X. Our approach leverages deep learning-based structural prediction, incorporates standard physicochemical features of structural stability, and quantifies their contributions to a unified deleteriousness score to aid in downstream interpretation. When evaluated on a dataset of validated missense variants, AlphaRING-X outperformed gold-standard software in its classifications, while also attributing its predictions to the underlying structural characteristics. Finally, to facilitate broader use, AlphaRING-X has been made available as an open-source software package that, along with its modular and flexible design, enables adaptation and expansion of its implementation across differing use cases.

## Methods

### AlphaRING-X algorithm

#### Aims

For AlphaRING-X, we aimed to develop a highly accurate predictor of missense variant deleteriousness that retains interpretability and flexibility. Our approach focuses on comprehensively assessing how missense variants alter protein structural stability, as this is a key mechanism by which they contribute to phenotypic diversity [22,23]. To achieve this, we sought to capture structural stability at both local and global levels to minimise the loss of deleterious signal at either viewpoint. To retain interpretability, we aimed to capture these effects with as few features as possible and to translate them into a unified yet explainable deleteriousness score using a high-performance model whose feature contributions could be readily quantified. Finally, to maximise flexibility, we sought to release AlphaRING-X as open-source software that can predict from any user-provided amino acid sequence and single-residue substitution, rather than simply releasing a fixed set of predictions. Consistent with flexibility, AlphaRING-X is designed to use an in silico-derived feature space and to be modular and expandable, thereby avoiding input limitations caused by limited experimental data. This also allows for easy modifications to the tools employed to generate the feature space and to the feature space itself.

#### Feature space

Locally, a residue’s conformational flexibility and three-dimensional interactions are primary influences on its conservation and tolerance to substitution [24,25,26]. Due to this and their intuitive roles, we sought to capture the regional disorder and spatial adjacency of the substitution-site reference amino acid (SSRAA) within the AlphaRING-X feature space.

To capture the regional disorder, AlphaRING-X uses its first feature, the predicted local distance difference test (pLDDT) of the SSRAA. pLDDT is a per-residue confidence metric on local structure calculated by AlphaFold [27], which is based on the local distance difference test Cα [28] and is strongly correlated with structural order [29]. Therefore, pLDDT reflects AlphaFold’s confidence that a residue adopts a well-defined local three-dimensional environment, and that residues with high pLDDT are typically in ordered regions and residues with low pLDDT are typically in disordered regions. As a result, pLDDT offers a convenient estimate of regional disorder within an in silico-derived feature space, such as that of AlphaRING-X.

To capture spatial adjacency, AlphaRING-X uses residue interaction network (RIN) analysis. A RIN is a graph-based representation of a protein’s structure in which each residue is a graph node, and connections are drawn between nodes when pre-defined geometric criteria are met for specific covalent or non-covalent interactions, most typically interatomic distance cut-offs and angle constraints, thereby converting residue connectivity into graph edges. In RIN analysis, connectivity is summarised by degree, a per-residue metric equal to the total number of interaction edges formed by a residue node. Therefore, as its second feature, AlphaRING-X uses the degree of the SSRAA. To estimate degree, AlphaRING-X uses RING [30], the most widely used tool for RIN generation and analysis.

Globally, a widely used metric for measuring how a substitution impacts protein structural stability is the change in change of Gibbs free energy (ΔΔG). Therefore, AlphaRING-X uses the ΔΔG resulting from the substitution as its third feature. To predict ΔΔG, AlphaRING-X employs FoldX [31], a widely used tool that utilises an empirical force field to assess the energetic effect of mutations. While ΔΔG is an effective descriptor of global structural stability, it does not describe local structural stability, unlike pLDDT and degree. Therefore, these features serve to complement each other.

Finally, AlphaRING-X considers the location of the substitution within the protein by calculating the relative substitution position (RSP) as its fourth and final feature. RSP is a continuous value ranging from 0, which indicates the N-terminus, to 1, indicating the C-terminus.

#### Model

AlphaRING-X uses an efficient implementation of gradient boosting [32], XGBoost [33], to translate its feature space into a deleteriousness score. In gradient boosting, an XGBoost model uses an ensemble of decision trees, each of which defines a region-wise constant approximation to the mapping from features to deleteriousness via a hierarchy of simple decision rules. By iteratively adding trees that correct the residual errors of earlier trees and incorporating regularisation, along with built-in handling of missing values, XGBoost achieves strong predictive performance, surpassing even deep learning architectures on tabular data and on data with class imbalance [34,35], which are common in variant datasets, and present in AlphaRING-X’s training and test sets.

#### Prediction scoring

The AlphaRING-X model encodes the deleteriousness score as a probability, with higher values indicating greater deviation from neutrality. To classify a score as neutral or deleterious, AlphaRING-X follows the ACMG/AMP recommendation of having at least 90% certainty in its variant classification [4]. To achieve this, AlphaRING-X uses a lower threshold (LT), below which variants are classified as “neutral”, and an upper threshold (UT), over which variants are classified as “deleterious”. Scores between LT and UT are considered “ambiguous”. The LT and UT were derived from the maximum and minimum thresholds that achieved 90% classification precision for the neutral and deleterious classes respectively, in the AlphaRING-X test set. By using the maximum and minimum thresholds for LT and UT respectively, the probability range for an ambiguous score was minimised, increasing the likelihood that variants would be classified as neutral or deleterious, thereby aiding their downstream interpretation for clinical actionability.

#### Prediction interpretation

To ensure each deleteriousness score is interpretable, AlphaRING-X uses the Shapley additive explanations (SHAP) framework [36]. In this framework, each feature is assigned a SHAP value whose sign indicates whether the feature increases or decreases the deleteriousness score, and whose magnitude reflects the size of that effect. By design, the SHAP values for all features sum to the difference between the model’s score for a given variant and a baseline score, defined as the expected model output over a background set of variants. Here, SHAP uses the AlphaRING-X training set as background to calculate the baseline score.

#### Implementation

AlphaRING-X is implemented as a Linux-native and open-source Python package on GitHub [37] and archived on Zenodo [38], where detailed user instructions are available. In short, to predict the deleteriousness of missense variants, AlphaRING-X primarily requires two inputs: a reference amino acid sequence and one or more single-residue substitutions within that sequence (Fig. 1A).

**Fig. 1.**
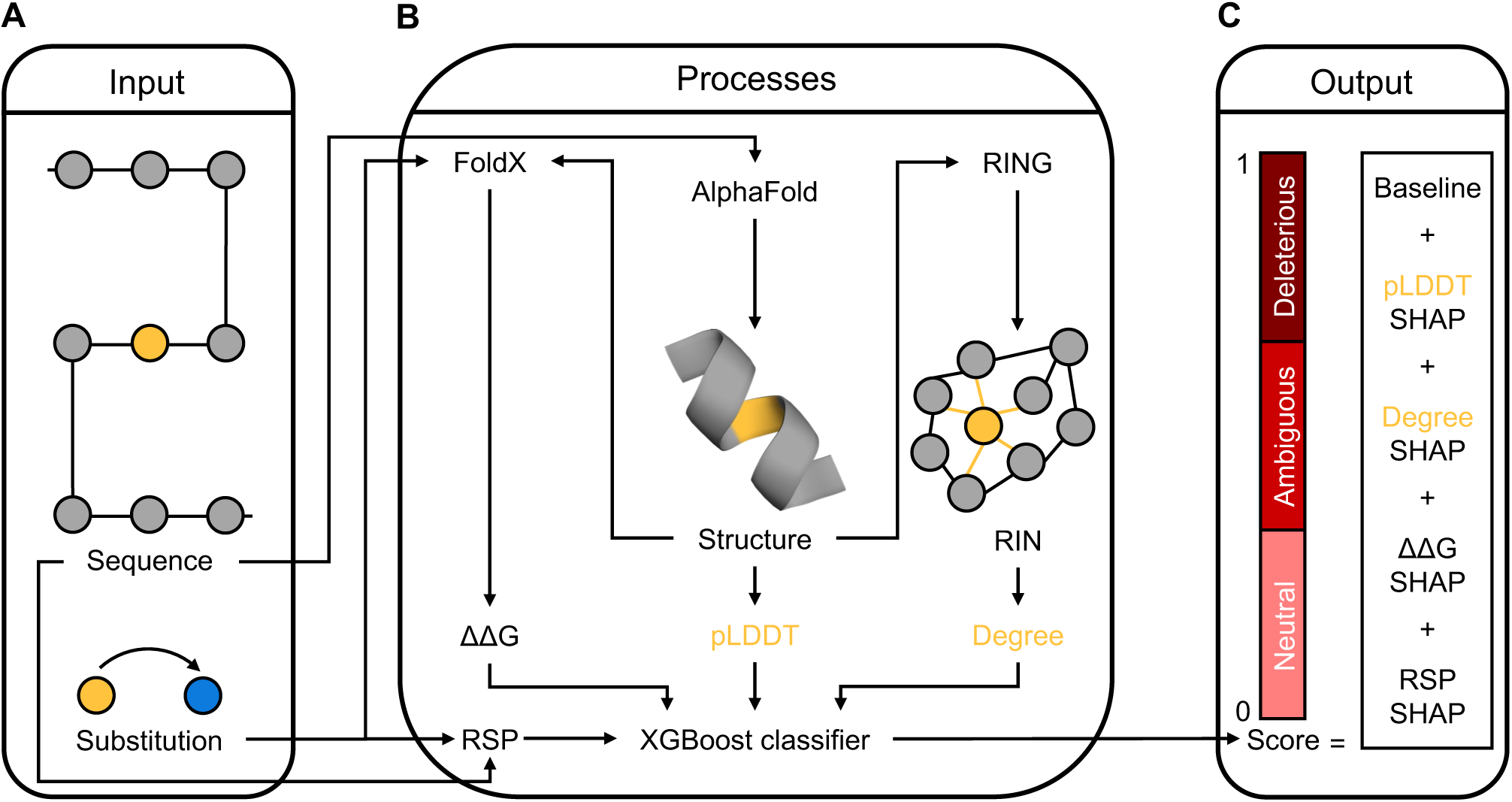
The AlphaRING-X algorithm. The reference and alternate amino acids at substitution sites are highlighted in yellow and blue, respectively, as are their derived properties. The remaining amino acids are in grey. Amino acids are either shown as circles or as part of an α-helix. The AlphaRING-X algorithm has been divided into three components: **A** inputs, **B** processes, and **C** outputs.

To process the inputs and extract features for prediction, AlphaRING-X performs the following steps: the reference sequence is modelled using AlphaFold [27] v2.3.2 to extract the pLDDT of the SSRAA; the model is converted into a RIN using RING [30] v4.0 to extract the degree of the SSRAA; the model is used to predict the ΔΔG caused by the substitution using FoldX [31] v5.1; and finally, the reference sequence and substitution are used to calculate the RSP, all of which are then passed to the XGBoost [33] v3.0.0 model (Fig. 1B). AlphaFold is run in monomer mode using its full database for multiple sequence alignment (MSA) and all available templates, but not accepting pre-computed MSAs. It requires a modern NVIDIA graphics processing unit to generate five models, relaxing its self-determined best model, which is passed to RING and FoldX. RING is run in relaxed mode to capture all possible interactions. FoldX is run using its PositionScan program.

Finally, the model accepts the extracted features and outputs a deleteriousness score as a probability, along with a “neutral”, “ambiguous”, or “deleterious” label, and quantifies the feature contributions to this score using SHAP [36] (Fig. 1C). For convenience, each substitution’s deleteriousness score, classification, feature values, and SHAP values are stored in a tabular format.

### Model development

All the datasets and scripts used for model development have been archived on Zenodo [39].

#### Dataset assembly

To assemble a gold-standard dataset for training the AlphaRING-X model, we retrieved missense variants from ClinVar [7,8] (accessed on 18^th^ February 2025). The variants were filtered to select single-nucleotide variants from the GRCh38 assembly that were determined as definitively “benign” (neutral) or “pathogenic” (deleterious) and marked as “reviewed by expert panel”. Additionally, residue-level variant deduplication was performed. This resulted in 1556 variants, of which 516 were classified as neutral and 1040 as deleterious. The feature values for these variants were calculated from the corresponding components of AlphaRING-X, yielding a tabular dataset.

#### Training

The tabular dataset was partitioned into an 80% training set and a 20% test set, stratified by class and grouped at the protein level, thereby preserving class proportions and ensuring that variants of the same protein did not appear in both sets. Next, a random search over XGBoost hyperparameters was performed to generate 150 combinations. Each hyperparameter combination was assessed by further splitting the training set into five folds using stratified and grouped K-fold cross-validation, where each fold was trained with weights to equalise the cost of errors across both classes. The combination with the highest mean area under the receiver operator curve (auROC) across the five folds was selected and refit on the whole training set. Next, the fully trained model was calibrated using cross-validation with sigmoid calibration, employing the same folds and weights as before.

Finally, the calibrated model was packaged with the whole training set, serving as the final model and the SHAP background dataset of AlphaRING-X respectively.

### Benchmarking

All datasets and scripts used for benchmarking have been archived on Zenodo [39].

#### Classification

To test the classification performance for each predictor of deleteriousness in this study, a mean auROC with standard deviation was calculated by conducting bootstrap resampling 2000 times on the test set. To compare the difference in predictor auROCs, the procedure of Cheng et al. [21] was followed, in which the proportion of the 2000 bootstrap resamples for which predictor A’s auROC was less than or equal to predictor B’s was calculated as a one-sided *p* value to represent the probability that the observed improvement would arise by chance under the null hypothesis that predictor A is not better than predictor B.

#### Resolution

For each predictor, target precision between 80% to 100% was assessed at 0.5% increments on the test set, and, at each target, an LT was set as the largest AlphaRING-X score meeting the precision level for the neutral class and a UT was set as the smallest score meeting the precision level for the deleterious class. Variants with scores ≤LT were classified as “neutral” (resolved), variants with scores ≥UT were classified as “deleterious” (resolved), and the remainder were considered “ambiguous” (unresolved).

## Results

### Benchmarking AlphaRING-X classification performance

To test AlphaRING-X on its ability to classify missense variants, we evaluated it on our test set comprising validated neutral and deleterious missense variants from ClinVar [7,8]. In doing so, we benchmarked AlphaRING-X against CADD [16], given the position of CADD as a widely adopted gold standard for variant deleteriousness prediction, and its complete prediction coverage of our test set. Instead of using logistic regression to make predictions, as CADD does [16], AlphaRING-X uses an XGBoost [33] model, a gradient-boosting [32] algorithm that often outperforms even deep learning architectures on tabular datasets and datasets with class imbalance [34,35], similar to those used to train and test AlphaRING-X. Consistent with this, testing showed AlphaRING-X achieved an auROC of 0.940 compared to CADD’s 0.895 (Fig. 2A), a significant difference (*p* < 0.05; bootstrap), demonstrating AlphaRING-X as a highly accurate discriminator between neutral and deleterious missense variants.

**Fig. 2.**
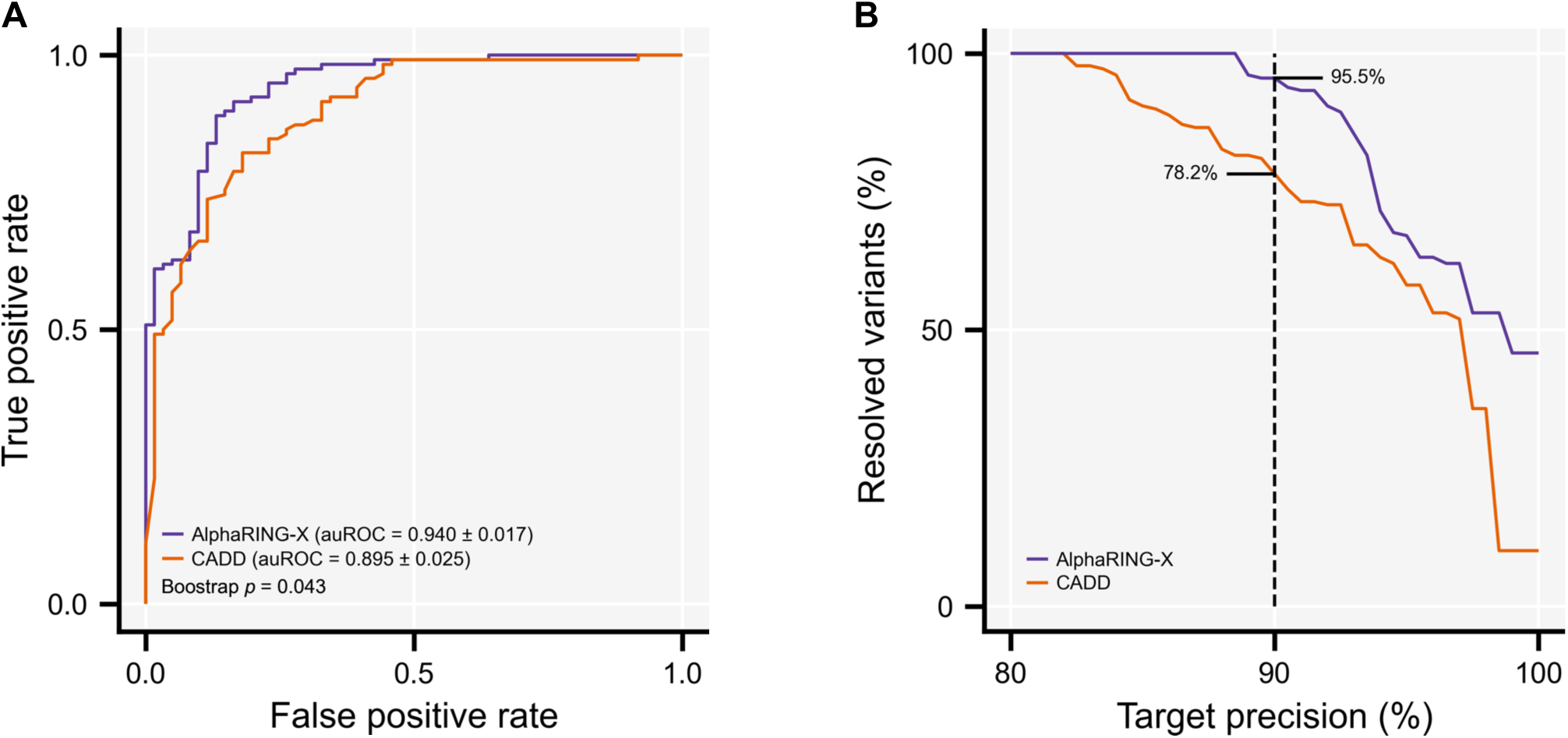
Performance of AlphaRING-X in missense variant classification. **A** Receiver operating characteristic analysis using the AlphaRING-X test set. Predictor auROCs are shown as the mean ± standard deviation of 2000 bootstrap resamples, alongside the one-sided *p* value for the difference in bootstrap-sampled auROCs between the predictors. **B** Percentage of resolved (unambiguously classified) missense variants at increasing target precision using the test set. Any given target precision required the precision level to be achieved on both the neutral and deleterious classes. The annotated dashed line denotes the percentage of resolved variants by each predictor at 90% precision.

For clinical applications, predictors must meet stringent target precision standards, with the ACMG/AMP recommending at least 90% certainty in variant classifications [4]. To that end, for both AlphaRING-X and CADD, we swept target precision from 80% to 100% on both neutral and deleterious variant classes. At all target precisions above 82.5%, AlphaRING-X was able to resolve (unambiguously classify) more variants than CADD. At 90% precision AlphaRING-X achieved 95.5% compared to CADD’s 78.2% (Fig. 2B), demonstrating AlphaRING-X as a more clinically actionable predictor of missense variant deleteriousness.

### Interpreting AlphaRING-X predictions

Besides being accurate, deleteriousness scores must also be explainable to support their downstream mechanistic interpretation and enhance their clinical utility. However, currently available predictors of missense variant deleteriousness do not provide this explainability on a prediction-by-prediction basis. To address this, AlphaRING-X uses SHAP [36], a framework that allows it to quantify the magnitude and direction of each feature’s contribution to every deleteriousness score.

We leveraged the use of SHAP in AlphaRING-X by using both absolute and signed SHAP values from our test set to draw broader conclusions about the importance of each feature (Fig. 3A) and its relationship with its corresponding SHAP values (Fig. 3B). We observed that the degree and pLDDT of the SSRAA were the most important features, with median absolute SHAP values of 0.128 and 0.103 respectively, reflecting the leading roles that local connectivity and disorder play in a residue’s tolerance to substitution [24,25,26]. Across the test set, the contribution from degree was symmetric, as lower degrees produced more negative SHAP values, pushing predictions towards neutrality, whereas higher degrees produced more positive SHAP values, pushing predictions towards deleteriousness. Therefore, substitutions of residues with fewer predicted interactions were less likely to lead to deleterious predictions, and vice versa. However, for pLDDT, the contribution was less symmetric, with a slight skew towards neutrality. Therefore, while low pLDDT values were more likely to push variants towards neutral predictions, high pLDDT values were less informative as drivers of deleterious predictions. After degree and pLDDT, the next most important feature was ΔΔG, with a median absolute SHAP value of 0.085. ΔΔG captured certain instances of global structure destabilisation upon substitution, driving these predictions strongly towards the deleterious and thus contributing less towards the neutral. Finally, RSP was the least important feature, with an absolute median SHAP value of 0.037. RSP best captured instances in which substitutions occurred at either end of the protein, with those near the N- and C-termini more likely to be deleterious and neutral predictions respectively.

**Fig. 3.**
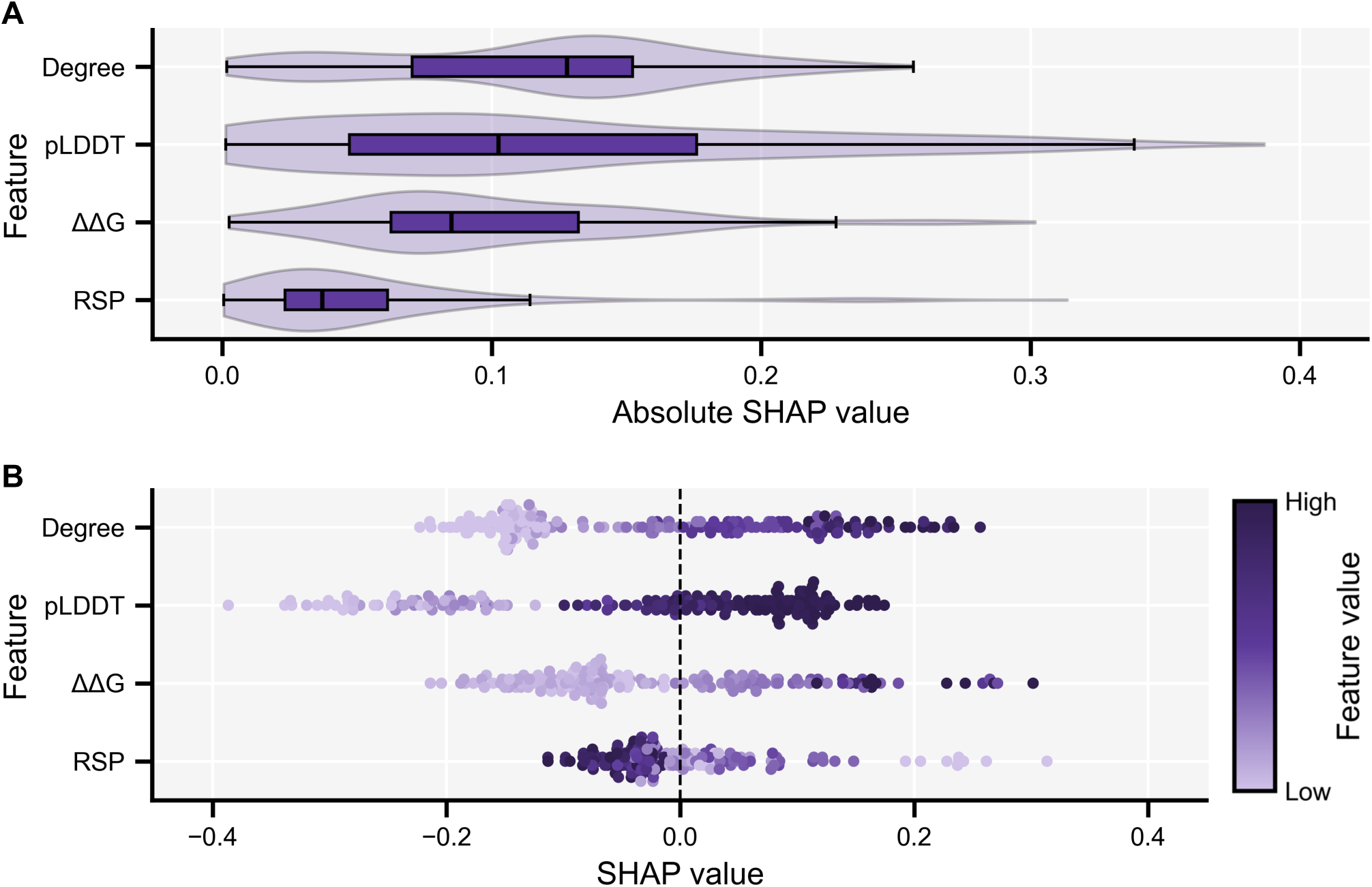
Contribution of features to AlphaRING-X predictions. **A** SHAP importance of each AlphaRING-X feature using the AlphaRING-X test set. Each feature is shown as a box plot overlaid by a violin plot representing that feature’s distribution of absolute SHAP values. **B** SHAP summary of each AlphaRING-X feature using the test set. Each feature is shown as a beeswarm plot, and each dot represents that feature’s signed SHAP value from an individual deleteriousness score.

After determining the relationship between each feature and its own SHAP values, we sought to characterise the relationships between the features. Therefore, we performed pairwise Spearman’s rank correlations (Additional File 1: Table. S1). The correlation between degree and pLDDT was the strongest and most significant (*r_s_* = 0.639, *p* = 6.01E-22), reflecting the fact that low and high values of both features tended to push predictions towards neutrality and deleteriousness respectively. However, the imperfection of this correlation was also intuitive, as, unlike degree, pLDDT contribution was not entirely symmetric, with its skew towards neutral outcomes. Therefore, while both features captured similar signals, they captured discrete information on deleteriousness. ΔΔG exhibited similar, albeit slightly less strong correlations with degree (*r_s_* = 0.401, *p* = 2.67E-08) and pLDDT (*r_s_* = 0.302, *p* = 3.88E-05), reflecting its shared yet reduced tendency to push predictions towards neutrality and deleteriousness. However, as with pLDDT, ΔΔG contribution was slightly skewed, but towards deleterious outcomes instead, suggesting it also captured discrete information. Finally, the correlations of RSP with the other features were the weakest and least significant (*r_s_* < 0.200, *p* > 0.01). While this was consistent with RSP’s observed auxiliary role in AlphaRING-X predictions, it also showed that RSP captured information orthogonal to that caught by the others.

### SHAP explains AlphaRING-X improvements over CADD

Providing interpretation towards AlphaRING-X predictions was necessary for its transparency and clinical utility. To maintain this transparency, we also sought to identify and explain the improvements of AlphaRING-X over CADD. Initially, we assessed the relationship between AlphaRING-X and CADD on our test set, using 90% precision thresholds for each: for AlphaRING-X, the LT and UT were 0.227 and 0.274 respectively, and for CADD, the LT and UT were 19.77 and 25.30 respectively. This showed a strong Spearman’s rank correlation (*r_s_* = 0.570, *p* = 1.09E-16), albeit with concordance of classification at only 67%. The remainder showed discordance or uncertainty, the latter caused by one or both predictors making an ambiguous call (Fig. 4A). To inspect the differences in classification further, we stratified the test set variants by whether their AlphaRING-X and CADD classifications matched their ClinVar classifications. Compared to CADD, AlphaRING-X was able to class match 11 more neutral and 16 more deleterious variants (Fig. 4B), representing 18% and 14% of the two variant classes respectively. This further confirmed the improvement of AlphaRING-X over CADD in classifying missense variants, but also revealed that this improvement was pronounced for both the neutral and deleterious classes.

**Fig. 4.**
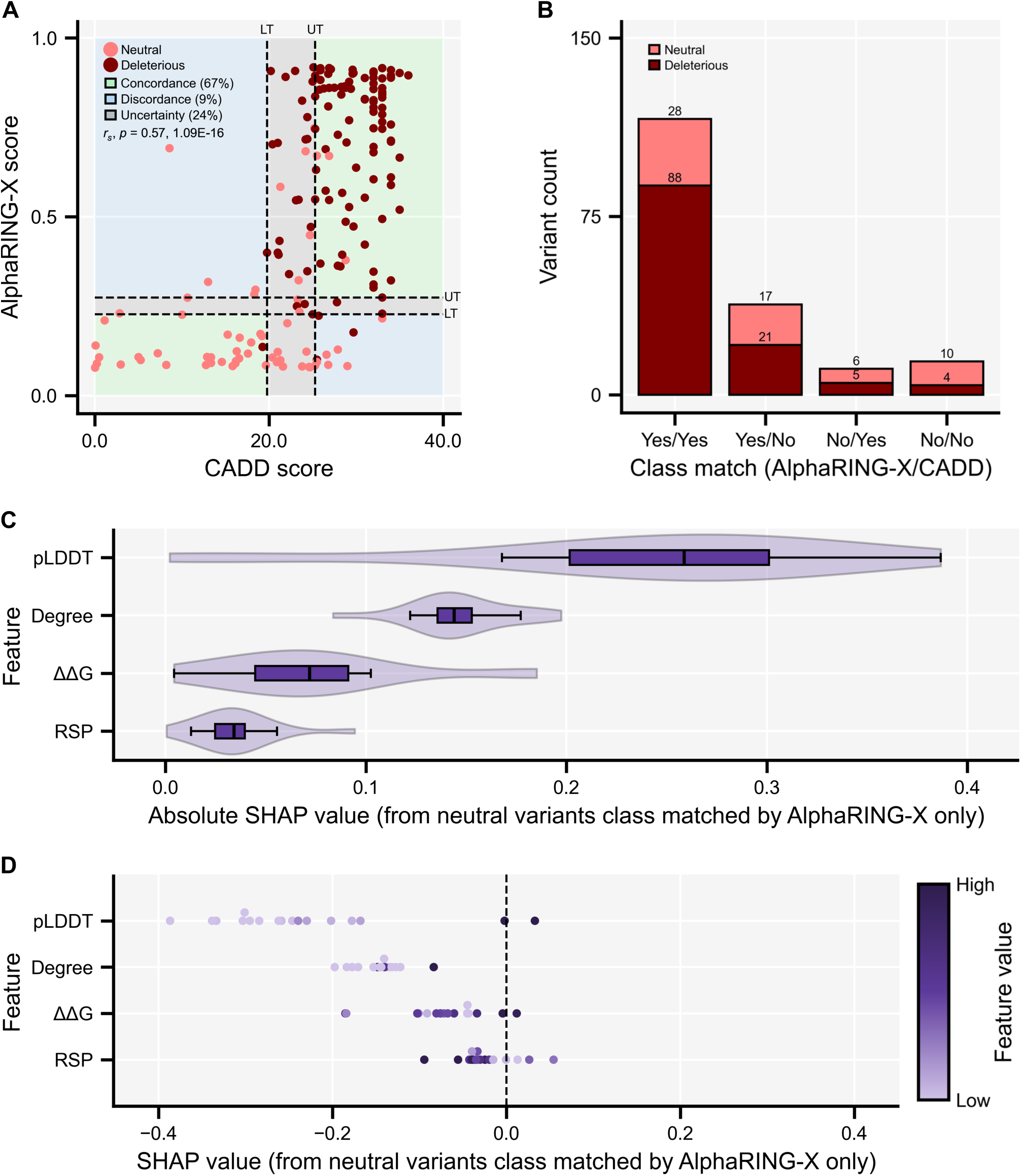
Explanation of the improvements of AlphaRING-X over CADD. **A** Scatterplot of AlphaRING-X versus CADD scores from the AlphaRING-X test set. LTs and UTs for AlphaRING-X and CADD are denoted by horizontal and vertical dashed lines respectively, showing class concordance, discordance, and uncertainty. Spearman’s *r_s_* and *p* value are shown. **B** Missense variants from the test set stratified by whether AlphaRING-X and CADD matched their classes on ClinVar. **C** SHAP importance of each AlphaRING-X feature for neutral variants classified correctly only by AlphaRING-X. Each feature is shown as a box plot overlaid by a violin plot representing that feature’s distribution of absolute SHAP values. **D** SHAP summary of each AlphaRING-X feature using the same test subset. Each feature is shown as a beeswarm plot, and each dot represents that feature’s signed SHAP value from an individual prediction.

Next, we stratified the absolute and signed SHAP values from the test set to isolate subsets of neutral and deleterious variants that were class matched by AlphaRING-X but not by CADD. This allowed us to investigate whether AlphaRING-X features contributed differently to the two subsets. Interestingly, the neutral subset showed the most pronounced change compared to the whole test set. Despite the importance of degree, ΔΔG, and RSP remaining similar, pLDDT became the most important feature by far, with a median absolute SHAP value of 0.259 (Fig. 4C). Factoring in SHAP directionality, it was clear that almost all of the neutral subset harboured predictions with very low pLDDT values and highly negative SHAP values (Fig. 4D). These pLDDT SHAP values also became negatively correlated with the other features, albeit with weaker statistical significance (Additional File 2: Table. S2), likely due to limited power from the relatively small subset size. Considering these changes, the improvement of AlphaRING-X over CADD on the neutral subset stemmed from the framework’s ability to capture how substitutions in residues with very low pLDDT (low order) result in a neutral outcome, with strongly negative SHAP values driving these correct neutral predictions.

Regarding the subset of deleterious variants class matched by AlphaRING-X but not by CADD, fewer changes in feature importance were observed compared to the whole test set (Additional File 3: Fig. S1A; Additional File 3: Fig. S1B). The pairwise Spearman’s rank correlations between SHAP values across different features reflected weaker versions of those from the whole set, except for those involving RSP (Additional File 4: Table. S3), which appeared more negative due to its wide but sparse spread of SHAP values in the positive direction. Therefore, the feature contributions towards AlphaRING-X predictions in the deleterious subset are consistent with those in the entire set.

## Discussion

In AlphaRING-X, we have developed the first predictor of missense variant deleteriousness that is both highly accurate and easily interpretable. We designed a feature space based on protein structural stability that captures changes at the local and global levels via widely used in silico tools, translating this into an interpretable deleterious score using explainable machine learning that quantifies each feature’s contribution. We have facilitated broader use and further development of AlphaRING-X by making it an open-source software package and by providing the datasets and scripts used for model development and performance benchmarking, enabling its adaptation and expansion.

Our performance benchmarking demonstrated that AlphaRING-X is a clinically viable option for the prediction of missense variant deleteriousness, as it not only outperformed a gold-standard tool in CADD [16] but was also able to unambiguously classify more variants as neutral or deleterious at clinically relevant precisions set out by guidelines such as those from the ACMG/AMP [4]. To maximise classification certainty, we developed and evaluated the AlphaRING-X model using stringent ClinVar criteria. This meant that, unlike other methods, we only considered missense variants reviewed by an expert panel and opted not to include variants with moderate levels of support. Although this decision likely reduced noise around the ground-truth labels, enabling AlphaRING-X to capture a more accurate signal of deleteriousness during training, it also limited the dataset size for model development and evaluation, likely reducing the diversity of variants seen by AlphaRING-X. Therefore, further development of AlphaRING-X should include larger, high-quality datasets by aggregating clinically reviewed variants from ClinVar and other reliable sources, such as ClinGen [5,6] and COSMIC [9].

Furthermore, benchmarking AlphaRING-X using this high-quality dataset limited our ability to compare it with other tools because of their incomplete prediction coverage of the dataset. This was especially clear with AlphaMissense [21], which released a highly accurate but incomplete set of predictions, without providing its model for users to make new predictions. This highlights the need for open-source frameworks to support innovative benchmarking efforts. However, datasets of variants, such as those in ProteinGym [40], have been developed for standardised benchmarking and, as a result, have been widely used to compare variant-effect predictors, including those mentioned in this study. Therefore, future work would also benefit from evaluating AlphaRING-X on these public datasets.

In addition to accuracy, our benchmarking also analysed the SHAP-based interpretation of AlphaRING-X predictions. This revealed that features describing the local structural stability of the SSRAA, namely degree and pLDDT, which capture local connectivity and disorder, made the strongest contributions, encoding related yet distinct information on deleteriousness. The importance of local features was even more pronounced for gold-standard neutral variants, which tend to be misclassified by other methods. Given the observed success of the local features, we believe that future efforts to extend the AlphaRING-X feature space should consider additional local structural characteristics. In particular, local packing [41,42] and solvent accessibility [43,44] are also regarded as primary influences on a residue’s tolerance to substitution, and they should be prioritised.

Regarding the feature space, AlphaRING-X was designed to thoroughly examine structural stability at both local and global levels, as changes in this are a key mechanism through which missense variants influence phenotypic diversity and disease [22,23]. However, this is not the only mechanism, as some variants affect other aspects of proteins, such as catalytic activity, binding, or expression. This has been thoroughly catalogued through the development of deep mutational scanning assays [45,46], a method that links selection for protein function with high-throughput DNA sequencing to measure the activity of protein variants at scale, thereby facilitating the generation of diverse datasets that assess the different mechanisms by which variants can impact proteins. These datasets have been compiled by sources such as ProteinGym [40] to assess the performance of variant-effect predictors across different mechanisms of variant impact, revealing varied results depending on each predictor’s design. Therefore, AlphaRING-X likely loses the deleteriousness signal from missense variants that affect proteins through mechanisms other than structural stability, and we suggest that future development of AlphaRING-X incorporate descriptors of these mechanisms into the feature space to maximise performance across missense variant types.

To facilitate the aforementioned adaptations to AlphaRING-X, it has been fully open-sourced [37,38,39], thereby enabling its implementation to be customised. In addition to modifying the feature space, users can also adjust or replace the in silico tools used to generate and translate it. The most consequential of these tools for generating the feature space is AlphaFold [27], which predicts protein structure from the reference amino acid sequence, directly providing the pLDDT feature and, indirectly, the degree and ΔΔG features. While AlphaFold is highly accurate at predicting monomeric protein structures, it requires a Linux machine and a ∼2.62 terabyte database to generate MSAs for its predictions, which often take multiple hours to complete. As a result, AlphaRING-X inherits the same computational limitations. To minimise these problems, AlphaRING-X checks whether a model has already been generated for the provided reference sequence, thereby scaling well when predicting multiple variants of the same protein. However, when predicting variants of different proteins, we believe users would greatly benefit from having a choice of protein structure prediction tools. In particular, ESMFold [47] would complement the current AlphaRING-X framework, as a marginally less accurate but less computationally demanding option. ESMFold is largely operating system-independent and does not use MSAs or an extensive database to generate them, cutting prediction time from hours to minutes. Importantly, like AlphaFold, ESMFold also computes pLDDT values, facilitating its integration into AlphaRING-X.

Finally, beyond improving the interpretation of human missense variants, AlphaRING-X’s framework has the potential to be adapted for non-human variants. Adapting AlphaRING-X for multi-species use would provide a unified framework for variant prioritisation in model organisms, agricultural species, and pathogens, where curated variant resources are relatively limited but high-quality protein sequences are readily available. Like other multi-species predictors [48,49,50,51,52], AlphaRING-X operates at the protein sequence level, predicting the deleteriousness of a missense variant from a single user-provided reference sequence and the substitution within it. Furthermore, the tools employed by AlphaRING-X and the feature space itself are fundamentally species-agnostic and independent of genomic coordinates or assemblies. To translate this potential into practice, expanding the model beyond its current human-centric training data would ensure robustness across phylogenetically diverse species. It is with this adaptation in mind that we opted to use the species-agnostic terms “neutral” and “deleterious” to describe variants, instead of “benign” and “pathogenic”.

## Conclusions

In conclusion, AlphaRING-X broadens access to accurate interpretation of missense variant deleteriousness, outperforming gold-standard tools while meeting the precision targets required by clinical guidelines. By coupling XGBoost with SHAP-based explanations of feature contributions, it delivers interpretable predictions that highlight local connectivity and disorder as key determinants of a residue’s tolerance to substitution. We expect that the open-source release of AlphaRING-X and its training and evaluation resources will aid the identification of clinically actionable variants in both research and diagnostic contexts and support the adaptation and expansion of the framework.

## List of abbreviations

ΔΔG: change in change of Gibbs free energy
ACMG/AMP: The American College of Medical Genetics and Genomics and the Association for Molecular Pathology
auROC: area under the receiver operator curve
LT: lower threshold
MSA: multiple sequence alignment
pLDDT: predicted local distance difference test
RIN: residue interaction network
RSP: relative substitution position
SHAP: Shapley additive explanations
SSRAA: substitution site reference amino acid
UT: upper threshold.

## Declarations

### Ethics approval and consent to participate

Not applicable.

### Consent for publication

Not applicable.

### Availability of data and materials

The AlphaRING-X software package is available at GitHub [37] and archived in Zenodo [38] for long-term accessibility. Additionally, the datasets and scripts used for model development and benchmarking are also archived in Zenodo [39]. All other data, including supplementary data, are available at bioRxiv online.

### Authors’ contributions

Conceptualisation: A.M.-L., E.B. Data curation: A.M.-L. Formal analysis: A.M.-L., E.B. Funding acquisition: A.M.-L., A.B., G.N.M. Investigation: A.M.-L., E.B. Resources: A.M.-L., R.S., A.B., G.N.M., E.B. Methodology: A.M.-L., R.S., A.B., G.N.M., E.B. Project administration: A.M.-L., G.N.M., E.B. Software: A.M.-L. Supervision: A.M.-L., G.N.M., E.B. Validation: A.M.-L. Visualisation: A.M.-L. Writing – original draft: A.M.-L. Writing – review & editing: all authors. All authors read and approved the final manuscript.

## Acknowledgements

Not applicable.

## Supplementary information

Additional File 1: Table. S1. Spearman’s rank correlation between each AlphaRING-X feature’s signed SHAP values using the AlphaRING-X test set.

Additional File 2: Table. S2. Spearman’s rank correlation between each AlphaRING-X feature’s signed SHAP values for neutral variants classified correctly only by AlphaRING-X.

Additional File 3: Fig. S1. Contribution of features to AlphaRING-X predictions for deleterious missense variants classified correctly only by AlphaRING-X. **A** SHAP importance of each AlphaRING-X feature for deleterious variants classified correctly only by AlphaRING-X. Each feature is shown as a box plot overlaid by a violin plot representing that feature’s distribution of absolute SHAP values. **B** SHAP summary of each AlphaRING-X feature using the same test subset. Each feature is shown as a beeswarm plot, and each dot represents that feature’s signed SHAP value from an individual deleteriousness score.

Additional File 4: Table. S3. Spearman’s rank correlation between each AlphaRING-X feature’s signed SHAP values for deleterious variants classified correctly only by AlphaRING-X.

