## Supplementary material for "AlphaRING-X: accurate interpretation of missense variant deleteriousness based on protein structural stability": Additional File 1.pdf

| Feature SHAP one | Feature SHAP two | $r_s$ | $p$ |
|---|---|---|---|
| Degree | pLDDT | 0.639 | 6.01E-22 |
| Degree | ΔΔG | 0.401 | 2.67E-08 |
| Degree | RSP | 0.011 | 0.884 |
| pLDDT | ΔΔG | 0.302 | 3.88E-05 |
| pLDDT | RSP | 0.032 | 0.674 |
| ΔΔG | RSP | 0.154 | 0.040 |
