## Supplementary material for "AlphaRING-X: accurate interpretation of missense variant deleteriousness based on protein structural stability": Additional File 2.pdf

| Feature SHAP one | Feature SHAP two | $r_s$ | $p$ |
|---|---|---|---|
| pLDDT | Degree | -0.441 | 0.076 |
| pLDDT | ΔΔG | -0.701 | 0.002 |
| pLDDT | RSP | -0.691 | 0.002 |
| Degree | ΔΔG | -0.007 | 0.978 |
| Degree | RSP | 0.044 | 0.866 |
| ΔΔG | RSP | 0.466 | 0.060 |
