## Supplementary material for "AlphaRING-X: accurate interpretation of missense variant deleteriousness based on protein structural stability": Additional File 4.pdf

| Feature SHAP one | Feature SHAP two | $r_s$ | $p$ |
|---|---|---|---|
| Degree | pLDDT | 0.606 | 0.004 |
| Degree | ΔΔG | 0.350 | 0.120 |
| Degree | RSP | -0.656 | 0.001 |
| pLDDT | ΔΔG | 0.203 | 0.377 |
| pLDDT | RSP | -0.632 | 0.002 |
| ΔΔG | RSP | 0.045 | 0.847 |
