## Supplementary figures and images for "AlphaRING-X: accurate interpretation of missense variant deleteriousness based on protein structural stability"

### Additional File 3.pdf

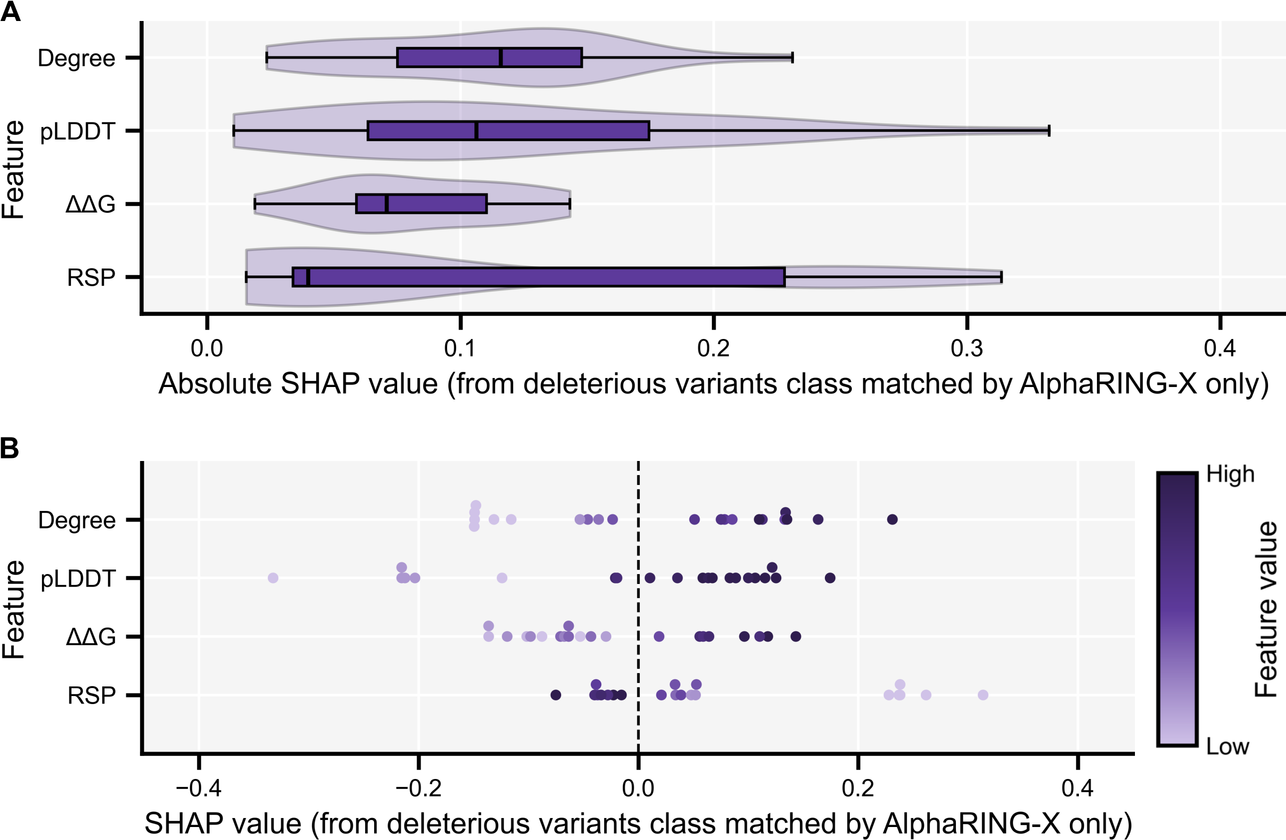
